# Preclinical Evaluation of [^18^F]Itareparib, a brain penetrant PARP1-selective radioligand, in Glioma Rat Model and Nonhuman Primate

**DOI:** 10.64898/2026.10.16.758050

**Authors:** Peng Wen Tan, Nira Hernandez-Martin, Claudia Esposito-Zapero, Christina Sun, Takuya Toyonaga, Ming-Qiang Zheng, Michael Kapinos, Monique A Thomas, Syed Faraz Nejati, Clement Korsah, Daniel Holden, Baosheng Chen, Jorge Sepulcre, Richard E. Carson, Yiyun Huang, Henk M. De Feyter, Zhengxin Cai

**Affiliations:** Department of Radiology and Biomedical Imaging, Yale School of Medicine, New Haven, Connecticut, USA; CIC biomaGUNE, Basque Research and Technology Alliance (BRTA), Donostia–San Sebastián, Spain; Department of Biomedical Engineering, Yale School of Engineering and Applied Science, New Haven, Connecticut, USA; Department of Pharmacology, Yale School of Medicine, New Haven, Connecticut, USA

**Keywords:** PARP1, PET, Itareparib, AZD9574, BT142, glioma

## Abstract

**Rationale:** Poly(ADP-ribose) polymerase-1 (PARP1) is a central mediator of DNA damage response and is upregulated in glioma, supporting it as a molecular imaging biomarker. However, quantitative PET assessment of PARP1 in infiltrative tumors with a preserved blood-brain barrier remains an unmet need. In this study, we evaluated if [^18^F]Itareparib, a brain-penetrant, non-trapping, PARP1-selective radioligand with reversible kinetics, could improve brain tumor diagnosis and enable target occupancy studies in neurooncology.

**Methods:** [^18^F]Itareparib was synthesized and evaluated in an orthotopic rat model of BT142 IDH1-mutant gliomas. Dynamic PET scans were performed with or without the administration of the PARP1-selective inhibitor AZD9574. Standardized uptake value ratios (SUVRs) for tumors were calculated using the contralateral region as reference and compared with distribution volume ratios (DVRs) calculated using kinetic modeling. PARP1-specific binding and tracer stability were assessed using autoradiography, immunofluorescence, biodistribution, and radiometabolite analysis. Brain kinetics of [^18^F]Itareparib were evaluated in a healthy nonhuman primate (NHP) at baseline and blocking conditions.

**Results:** [^18^F]Itareparib rapidly entered the rat brains and showed 4-fold higher uptake in BT142 tumors than in the contralateral brain. AZD9574 reduced tumor uptake by 77%, supporting the PARP1-specific binding of [^18^F]Itareparib in the tumors. SUVR_(30–60 min)_ showed the closest agreement with DVR. Autoradiographic tumor-to-reference mean ratio was 7.3, and AZD9574 reduced the tumor-to-reference ratio by 56%. PARP1 immunofluorescence was higher in tumor than the rest of the brain and correlated with autoradiographic results. In rats, the parent fraction of [^18^F]Itareparib exceeded 99% in brain through 60 min post tracer injection (*p.i.*) and was approximately 88% in plasma at 60 min *p.i.* In the NHP, [^18^F]Itareparib showed rapid brain uptake, and *V*_T_ decreased by 21 ± 6 % after blocking with AZD9574.

**Conclusion:** [^18^F]Itareparib showed brain penetration, high *in vivo* metabolic stability, reversible binding kinetics, and PARP1-specific tumor uptake. These findings support further development of [^18^F]Itareparib for the quantitative PET imaging of PARP1 in brain tumors.

## INTRODUCTION

Gliomas are diffusely infiltrative primary brain tumors associated with substantial morbidity and mortality. In clinical practice, magnetic resonance imaging (MRI) is the primary imaging modality for anatomical characterization of glioma and high-resolution assessment of tumor morphology (*1*). However, non-enhancing infiltrative tumor may extend beyond MRI-defined abnormalities, and treatment-related changes can be difficult to distinguish from recurrent disease (*2,3*). Molecular PET imaging can complement MRI by providing insight into tumor biology, with potential applications in diagnosis, prognosis, surgical planning and longitudinal disease assessment (*4*).

Poly(ADP-ribose) polymerase-1 (PARP1) is the most abundant and extensively studied member of the PARP family, owing in part to its central role in the DNA damage response (*5*). Following recognition of DNA single-strand breaks, PARP1 catalyzes poly(ADP-ribosyl)ation to recruit and coordinate proteins involved in base-excision repair (*6,7*). Replicative stress, genomic instability, and oxidative DNA damage are common features of glioma that contribute to the activation of DNA-damage signaling, making PARP1-mediated repair particularly relevant (*8*). Elevated PARP1 expression has been reported in human gliomas and has been implicated in tumor-cell survival and therapeutic resistance (*9–11*). These findings have supported the investigation of PARP inhibitors (PARPis) in glioma, including in combination with radiotherapy and temozolomide (*12–14*). The therapeutic interest in PARP1 further provides a rationale for the development of PARP1-targeted PET radioligands to assess PARP1 availability in glioma *in vivo* (*15–19*).

A PARP1-selective PET radiotracer could assess target availability beyond the sampled biopsy region and characterize regional variation within the tumor (*20–24*). For PARP1 PET imaging of infiltrative glioma, the radioligand should exhibit high affinity and selectivity for PARP1, sufficient brain penetration to access tumor regions with preserved blood-brain-barrier (BBB) integrity, low nonspecific retention in normal brain to achieve favorable tumor-to-background contrast, and reversible kinetics to enable the quantification of PARP1 (*25–27*). This motivated our recent development of [¹¹C]PyBic, a PARP1/2 PET tracer with excellent brain imaging properties (*26*). To extend these efforts to a fluorine-18-labeled PARP1 selective tracer, Itareparib, a non-trapping inhibitor, was selected as the scaffold for radiolabeling based on its excellent BBB penetration and high PARP1 binding affinity (*28*). Herein, we report the preclinical evaluation of [^18^F]Itareparib for quantitative imaging of PARP1 in a glioma rat model with a preserved BBB, and a healthy nonhuman primate (NHP).

## METHODS AND MATERIALS

### Chemistry and Radiochemistry

Detailed chemical synthesis and radiolabeling procedures are provided in Supplementary Information (S.I.).

### Measurement of lipophilicity (log D_7.4_)

Lipophilicity was determined using a modified method adapted from previously reported procedure (*29*).

### *Ex vivo* biodistribution

*Ex vivo* biodistribution was evaluated in wild-type Long-Evans rats (n = 25, 22 weeks, Charles Rivers Laboratories) to characterize the normal-organ distribution and clearance kinetics of [^18^F]Itareparib. [^18^F]Itareparib was administrated intravenously via the tail vein (approximately 10 MBq/rat) and subsequently euthanized at 5 min (n = 3), 10 min (n = 4), 15 min (n = 6), 30 min (n = 3), 60 min (n = 6), and 90 min (n = 3) post-injection (*p.i.*). For blocking studies, rats (n = 4) were administered with AZD9574 (0.5 mg/kg, *i.v*.) 15 min before [^18^F]Itareparib injection and euthanized at 15 min *p.i*. Blood was collected immediately before transcardial perfusion, after which the brain, lung, heart, liver, spleen, kidneys, muscle, and bone were harvested and weighed. Blood and tissue samples were quantified using an automated γ-counter (Wizard, PerkinElmer). A separate blood aliquot was collected for plasma radiometabolite analysis (see **Radiometabolite analysis** and S.I.). Each brain from rats euthanized at 15, 30, and 60 min *p.i.* was divided into two hemispheres: one hemisphere was used for biodistribution analysis, whereas the contralateral hemisphere was used for brain-homogenate radiometabolite analysis. Radioactivity concentrations were normalized to sample weight and injected dose, decay-corrected, and expressed as percentage injected dose per gram of tissue (%ID/g).

### BT142 animal model

Male adult RNU rats (Fox1RNU-Nude, 10–12 weeks old, Charles River Laboratories) were anesthetized with isoflurane using a 70/30 N₂O/O₂ carrier gas mixture and placed in a stereotactic rodent frame (David Kopf Instruments). An antiseptic treatment was provided before the scalp incision. A drill (0.6 mm bit) was used to perforate the skull at the coordinates corresponding to the allocation of the right hemisphere striatum according to the following bregma coordinates: AP 0.25 mm; LM 3.00 mm; DV 3.00 mm. BT142 tumor cells were injected using a 25 µL, 25 gauge-needle Hamilton syringe (Hamilton Company, USA). Bone wax (Ethicon) was used to cover the burr hole after suturing the incision with 3.0 suture thread (Monocryl Plus Antibacterial, Ethicon). Peri-and post-operative care included the use of analgesics (Lidocaine: <7 mg/kg, Meloxicam: 1–2 mg/kg, and Buprenorphine: 0.01–0.05 mg/kg). MRI was used to monitor tumor growth (See SI 1.5 for details). All animal procedures were approved by the Yale University Institutional Animal Care and Use Committee (*30*).

### PET imaging in BT142 rats

A total of 8 BT142-bearing rats underwent 90-min dynamic PET scans (baseline n = 5; blocking n = 3) on the FOCUS-220 scanner (Siemens, USA). [^18^F]Itareparib (23 ± 6 MBq in 400 μL, *i.v*.) was administered via the tail vein. For blocking studies, AZD9574 (0.5 mg/kg in saline, *i.v.*) (*31*) was injected via tail vein 10 min prior to the radiotracer injection. A 9-min transmission scan was acquired for attenuation correction right after the emission scan. Images were reconstructed using 3D ordered subset expectation maximization (OSEM3D; 2 iterations, 16 subsets) followed by a maximum a posteriori algorithm (MAP; 25 iterations), with corrections applied for decay, attenuation, scatter, normalization, and random events. Scans were reconstructed into 27 frames (6 × 30 s, 3 × 60 s, 2 × 120 s, and 16 × 300 s). Detailed analytical methods are described in S.I.

### *In vitro* autoradiography

*In vitro* autoradiography with [^18^F]Itareparib was performed on brain slices from a subset of the BT142 rats (n = 3, 2 slices per animal) used in the *in vivo* PET imaging. Baseline and blocking studies were conducted to assess its binding specificity. Brain tissues were sectioned into 20 µm slices axially and stored at -80 °C before use. Detailed methods are described in S.I.

### Immunofluorescence

Axial brain sections adjacent to those used for in vitro autoradiography from BT142 rats (n = 3; 2 sections per animal) were immunostained for PARP1 and human nuclear antigen (HuNu), imaged by fluorescence microscopy, and analyzed using FIJI-ImageJ, with detailed methods provided in the S.I.

### PET Imaging in NHP

Dynamic PET imaging was performed in a NHP (12 years old, female, 7.5 kg) under one baseline and two blocking conditions. [^18^F]Itareparib (184.7 ± 0.3 MBq) was injected intravenously with or without the blocking drug, AZD9574 (0.05 mg/kg or 0.15 mg/kg, *i.v*. 10 min before tracer injection) (*24,32*). Dynamic PET data were acquired for 120 min using a FOCUS 220 scanner and reconstructed by Fourier rebinning and filtered back projection with a Shepp–Logan filter, as previously described (*26*). Detailed analysis method is described in SI.

### Radiometabolite analysis

Blood and brain samples were collected from wild-type rats during *ex vivo* biodistribution studies at 15, 30, and 60 min *p.i*. Arterial blood samples were collected from NHP for gamma counting and radiometabolic analysis. Detailed analytical procedures are provided in S.I.

### Statistics

Statistical analyses were performed using a two-tailed unpaired student’s t-test or two-way ANOVA in GraphPad Prism 11. Simple linear regression analysis was employed to correlate PET parameters and autoradiography and immunofluorescence results. Data are presented as mean ± standard deviation (SD). A p-value of less than 0.05 was considered statistically significant.

## RESULTS

### Chemistry and Radiochemistry

[^18^F]Itareparib was synthesized with non-decay-corrected radiochemical yields of 3.43 ± 0.02%, molar activity of 126 ± 44 MBq/nmol, and radiochemical purity of greater than 98% (n = 4). The identity of the purified product was confirmed by coelution with Itareparib cold reference on radio-HPLC. The complete production process, from trapping of ^18^F^-^through synthesis, purification, and formulation, took approximately 120 min **(Scheme 1, Suppl. Figure 2 & 3)**.

**Scheme 1.**
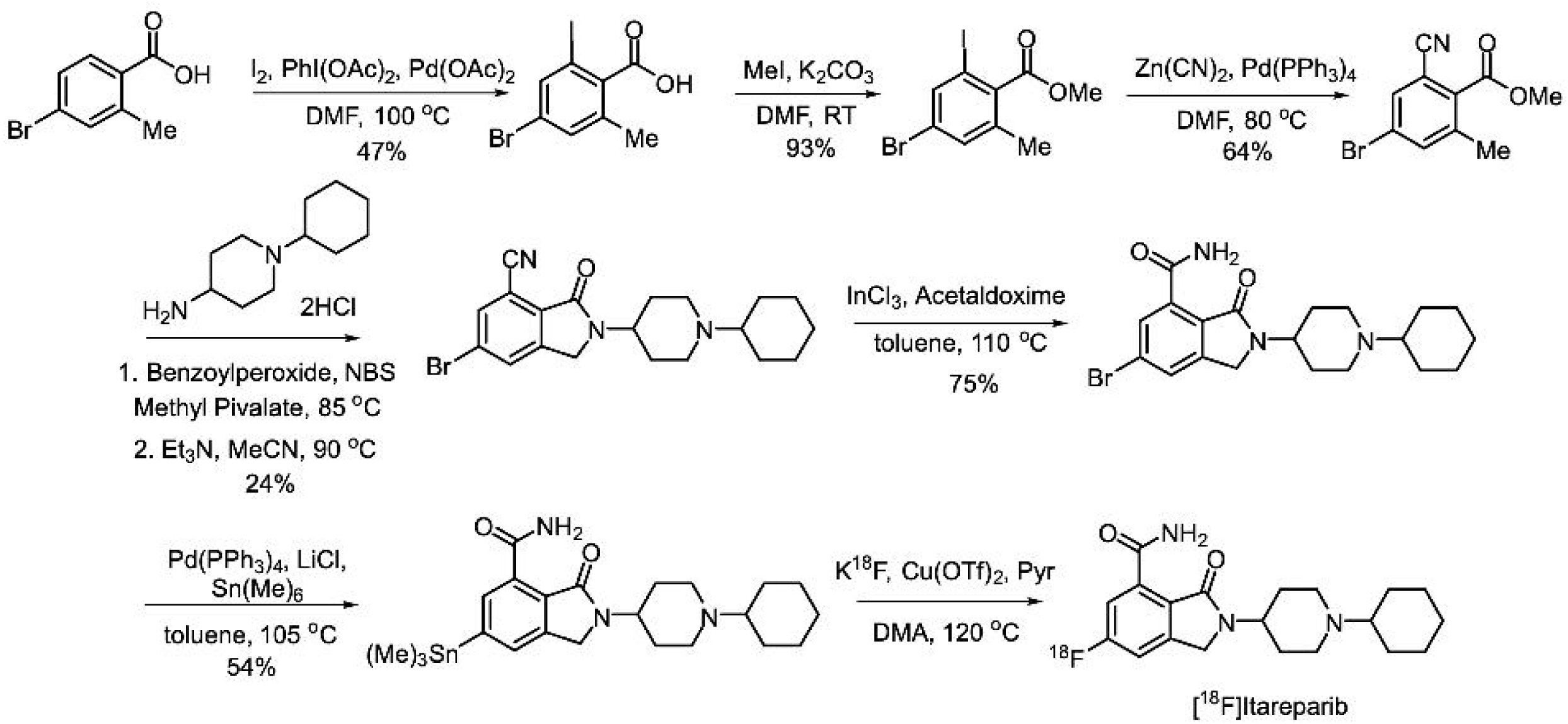
Synthetic route to the radiolabeling precursor and radiosynthesis of [¹⁸F]Itareparib.

### LogD Measurement

The measured LogD_7.4_ of [^18^F]Itareparib was 1.56 ± 0.04 (n = 4), within the optimal physicochemical properties for BBB penetration (*33*).

### *Ex vivo* biodistribution

*Ex vivo* biodistribution of [^18^F]Itareparib was assessed in healthy rats from 5 min to 90 min *p.i*. Brain uptake decreased from 0.139 ± 0.016 %ID/g at 5 min (n = 3) to 0.028 ± 0.005 %ID/g at 90 min (n = 3), accompanied by a reduction in blood radioactivity from 0.031 ± 0.003 %ID/g to 0.006 ± 0.001 %ID/g. Uptake in the heart, lungs, muscle, spleen, and bone also declined over time. Among the peripheral tissues examined, the liver and kidneys exhibited the highest activity concentrations. Hepatic uptake significantly increased at later time points and reached the peak at 60 min (2.090 ± 0.437 %ID/g, n = 6 vs 0.838 ± 0.197 %ID/g at 5 min *p.i.,* n = *3*, p < 0.001), whereas renal uptake peaked at 15 min (1.384 ± 0.562 %ID/g, n = 6 vs.865 ± 0.292 %ID/g at 5 min *p.i.*, n = 3, p = 0.009). The administration of the blocking drug resulted in a statistically significant decrease in spleen (0.559 ± 0.058 %ID/g, n = 4) in comparison to the baseline (0.963 ± 0.186 %ID/g, n = 6, p = 0.003) at 15 min *p.i*. Furthermore, a reduction in signal intensity was observed in the lung, bone, and muscle but did not reach statistical significance **(Suppl. Figure 4)**.

### Radiometabolite study in rats

Radio-HPLC analysis of brain homogenates showed no detectable radiometabolites of [^18^F]Itareparib at any time point, with > 99% of the detected radioactivity corresponding to the parent [^18^F]Itareparib, which eluted at approximately 8.9 min. Plasma analysis revealed a minor radiometabolite peak at approximately 0.6 min. The plasma parent fraction decreased over time (98%, 95%, and 88% at 15, 30, and 60 min *p.i*., respectively, **Suppl. Figure 5)**.

### Quantitative analysis of BT142 rats PET data

Dynamic PET imaging of BT142 tumor-bearing rats demonstrated rapid tumor uptake of [^18^F]Itareparib (**Figure 1 A**), reaching a peak SUV of approximately 1.8 within 5 min *p.i*. (**Figure 1 B)**. Under baseline conditions, tumor activity remained higher than the corresponding reference-brain region throughout the 90-min scan, while [^18^F]Itareparib was washed out of brain progressively. Tumor uptake of [^18^F]Itareparib was reduced by AZD9574 to the same level as the reference region after approximately 40 min *p.i.* (**Figure 1 B)**. Reference region activity peaked shortly after tracer administration and subsequently declined rapidly under both baseline and blocking conditions. Regional SUVRs calculated over the 30 – 60, 40 – 70, 50 – 80, and 60 – 90 min windows were plotted against SRTMC-derived DVRs, and 30-60 min window was identified as the appropriate static imaging window for simplified quantitative analysis **(Suppl. Figure, 6 A-B)**. [^18^F]Itareparib demonstrated significantly higher uptake in tumors (SUV: 0.97 ± 0.11, n = 5) than in the contralateral reference region (SUV: 0.20 ± 0.03, n = 5, p < 0.001) under baseline conditions **(Suppl. Figure 6 B)**. Under the blocking condition, the tumor SUV (0.23 ± 0.10, n = 3, p < 0.001) approached that in the reference region (0.13 ± 0.04, n = 3, p < 0.001). Consistent with these findings, SUVR_(30–60 min)_ of tumor decreased from 4.97 ± 0.73 (n = 5) at baseline to 1.71 ± 0.64 (n = 3, p < 0.001) under blocking conditions (**Figure 1 C)**. SRTMC analysis showed a similar extent of reduction in tumor DVR, which decreased from 4.16 ± 0.66 (n = 5) at baseline to 1.51 ± 0.25 (n = 3, p < 0.001) under blocking conditions (**Figure 1 D)**.

**Figure 1.**
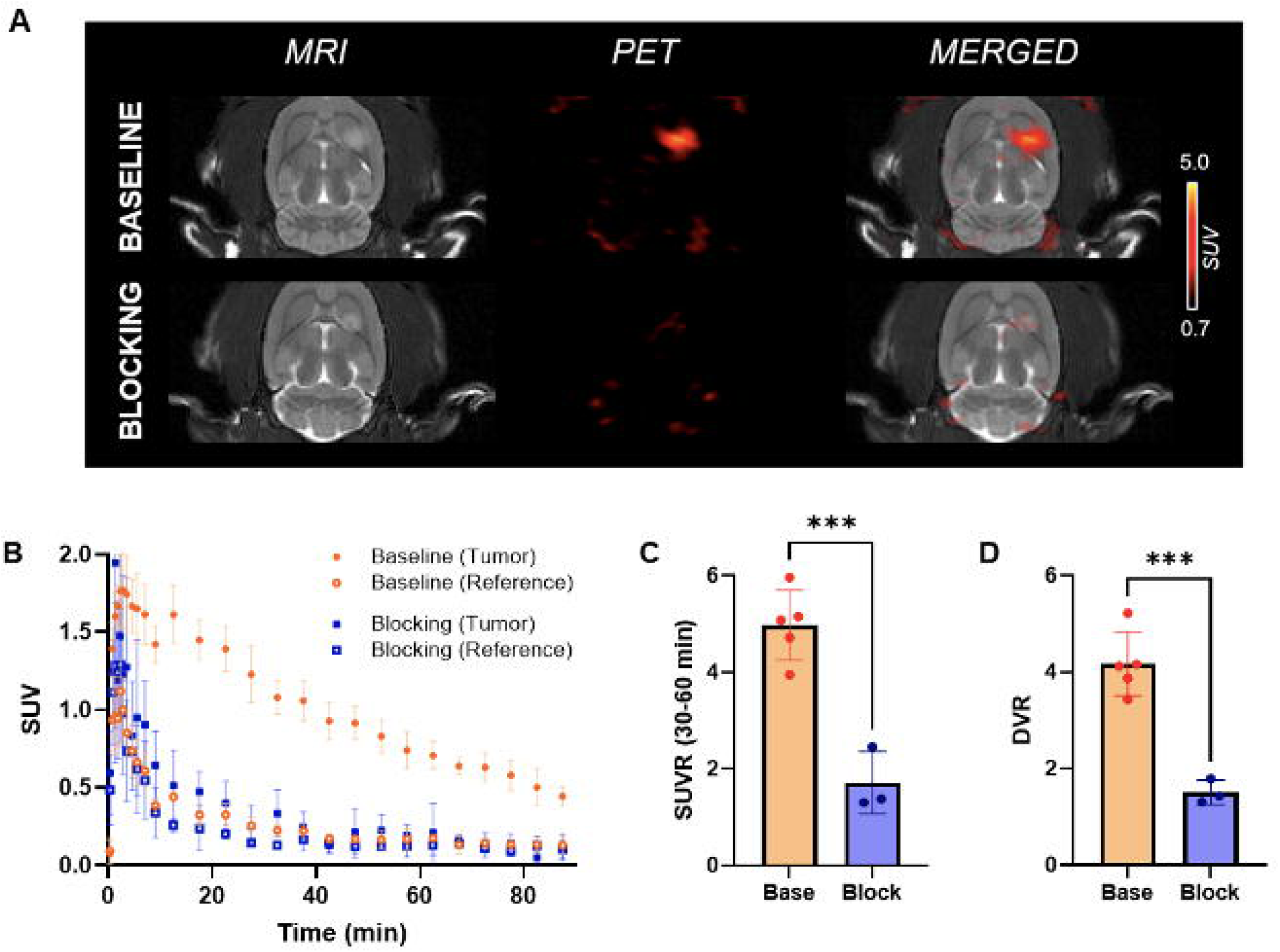
A) Representative MRI, PET and fused PET/MR images of [^18^F]Itareparib in two BT142 rats at baseline (upper row) and blocking conditions (bottom row); B) Mean SUV time-activity curves (TACs) of [^18^F]Itareparib in tumor and the contralateral non-tumoral region at baseline (orange, n = 5) and blocking conditions (blue, n = 3); C) Comparison of the tumor-to-reference SUVR_(30-60 min)_ of [^18^F]Itareparib in BT142 rats at baseline and blocking conditions. D) Comparison of the tumor-to-reference DVR of [^18^F]Itareparib in BT142 rats at baseline and blocking conditions. Student’s T-test. ***p < 0.001.

### *In vitro* autoradiography

*In vitro* autoradiography demonstrated greater [^18^F]Itareparib binding in BT142 tumor regions (n=3) than in the corresponding reference brain regions (**Figure 2 A**). Tumor uptake (540.3 ± 126.1 MBq ⋅ L⁻¹) was significantly higher than the reference region (98.3 ± 68.0 MBq ⋅ L⁻¹, p = 0.02). AZD9574 significantly reduced tumor uptake to 243.4 ± 164.4 MBq ⋅ L⁻¹ (n = 3, p = 0.03), while uptake in the reference region decreased by 57% to 40.9 ± 15.9 MBq ⋅ L⁻¹ (p = 0.41). The reduction in the reference region, although not statistically significant, was consistent with the decrease in tracer uptake observed in *in vivo* small-animal PET blocking studies (**Figure 2 B**). These findings corroborated the high tumor-to-reference contrast observed by PET in the BT142 glioma model.

**Figure 2.**
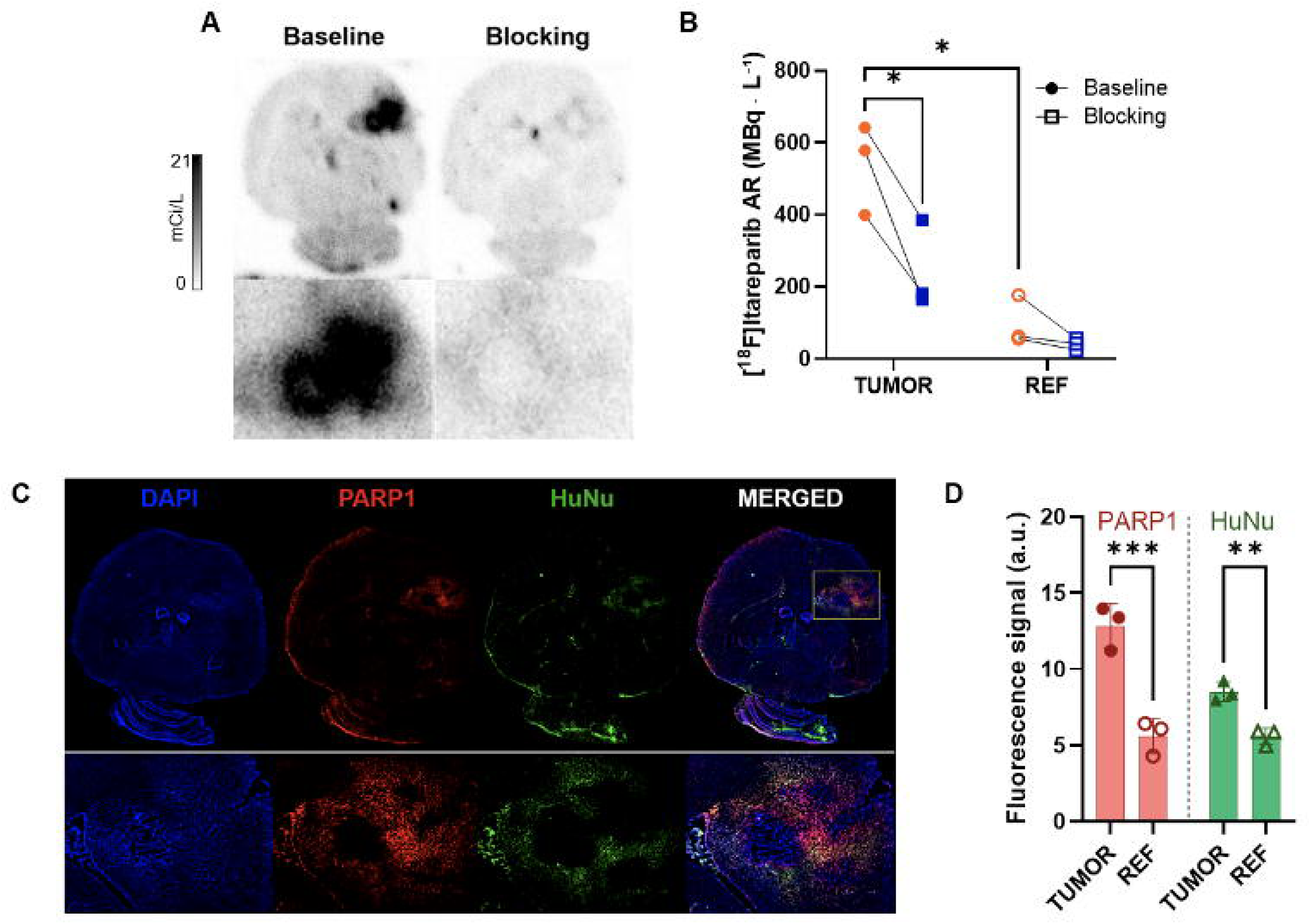
A) *In vitro* autoradiography (AR) of [^18^F]Itareparib with the BT142 rat brain sections at baseline (first column) and blocking conditions (second column); B) Comparisons of the AR signals of [^18^F]Itareparib in tumor and the contralateral non-tumor reference region at baseline (orange) and blocking conditions (blue); C) Representative images of immunofluorescence (IF) staining against PARP1 (red), HuNu (green) and DAPI (blue) in the BT142 rat brain sections; D) Quantitative analysis of the immunofluorescent signals of PARP1 and HuNu in tumor and the non-tumor reference region. Two-way ANOVA. *p < 0.05, **p < 0.01, ***p < 0.001.

### Immunofluorescence staining

Immunofluorescence analysis revealed increased PARP1 staining in HuNu-positive BT142 tumor regions compared with the contralateral brain region. The spatial correspondence between PARP1 and HuNu signals confirmed that the elevated PARP1 expression was associated with regions containing human tumor cells (**Figure 2 C)**. Quantitative analysis showed higher mean PARP1 fluorescence intensity (12.87 ± 1.45) and HuNu-positive staining (8.53 ± 0.65) in tumor ROIs than in contralateral ROIs (5.61 ± 1.14, n = 3, p < 0.001 and 5.6 ± 0.54, n = 3, p = 0.002 respectively, **Figure 2 D**). Both signals were correlated and heterogeneously distributed within the tumor (*R^2^*= 0.84, slope = 0.38, p = 0.01), consistent with the diffuse and infiltrative growth pattern of BT142 tumors **(Suppl. Figure 7 A)**. Both targets correlated with [^18^F]Itareparib autoradiography measures (PARP1: *R^2^* = 0.77, slope = 0.52, p = 0.02; HuNu: *R^2^*= 0.88, slope = 0.23, p = 0.01, **Suppl. Figure 7 B**).

### Radiometabolite study in NHP

The plasma free fraction (*f*_p_) of [^18^F]Itareparib in NHP plasma was 76 ± 4%. Radio-HPLC analysis of NHP plasma revealed radiometabolite peaks corresponding to those observed in rats, suggesting similar metabolic profiles across species. (**Figure 3 A**). The plasma parent fractions were 96% and 76% at 15 min and 90 min *p.i*., respectively, with an additional radiometabolite peak detected after 60 min *p.i*. Based on the absence of detectable radiometabolites in rat brain homogenates, NHP’s brain radioactivity was expected to predominantly represent parent [¹⁸F]Itareparib. The arterial input functions showed a rapid initial peak followed by a steep decline under both baseline and blocking conditions, with a higher initial peak in the blocking scan than in the baseline scan (**Figure 3 B)**.

**Figure 3.**
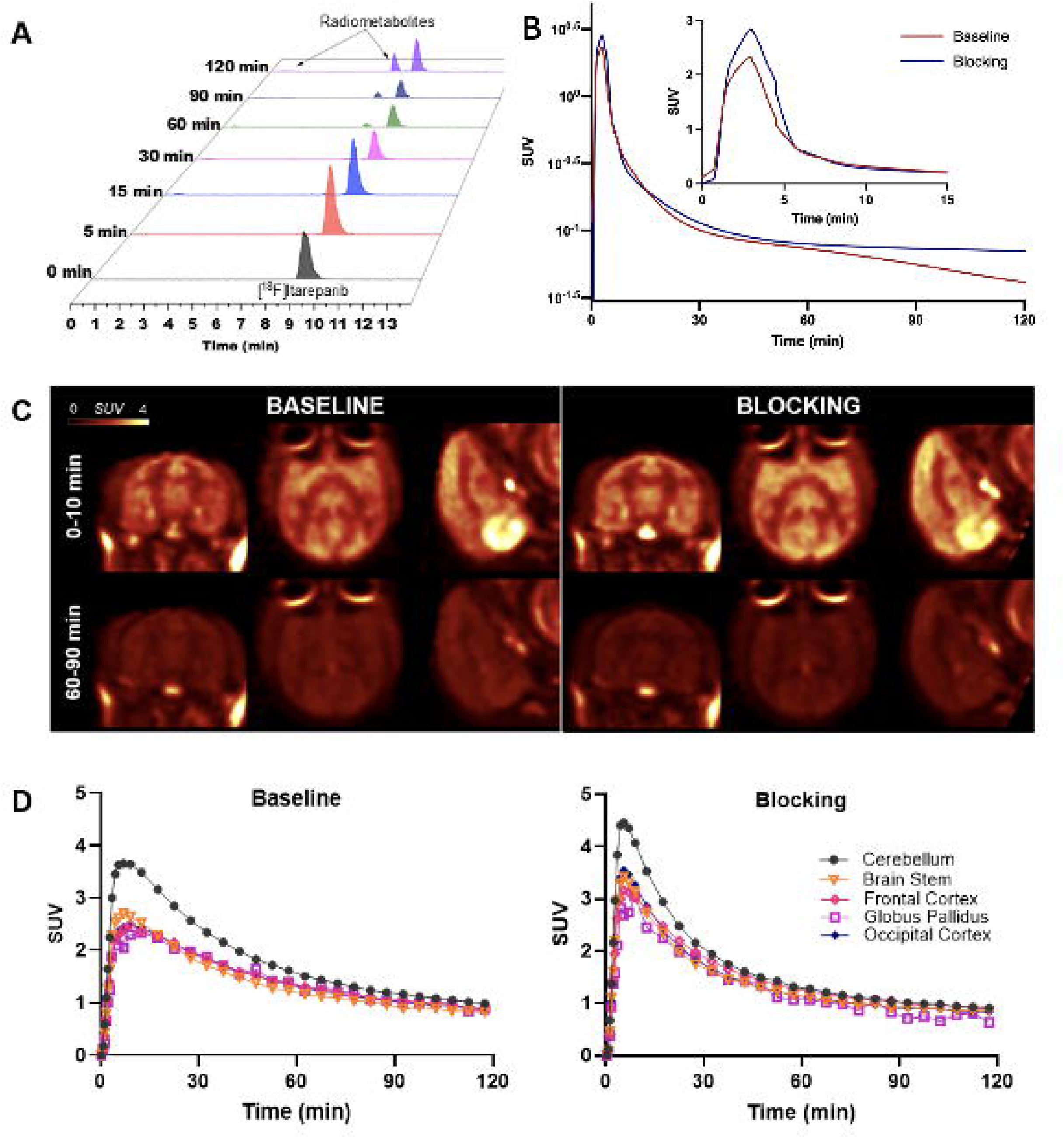
A) Radio-HPLC chromatograms of plasma samples from the baseline nonhuman primate (NHP) PET imaging study; B) Time course of the metabolite-corrected plasma concentration of [^18^F]Itareparib at baseline and blocking NHP PET scans; C) Summed SUV images of [^18^F]Itareparib in NHP brains from early (10-20 min *p.i*., top row) and later (60-90 min *p.i*., bottom row) time windows at baseline (left) and blocking (right) conditions; D) Time-activity curves (TACs) of [^18^F]Itareparib in selected brain regions of the NHP (cerebellum, brain stem, frontal cortex, globus pallidus, and occipital cortex) at baseline (left) and blocking (right, AZD9574, 0.05 mg/kg) conditions.

### NHP brain PET imaging

Dynamic NHP PET imaging demonstrated rapid brain penetration of [^18^F]Itareparib, peaking at 5 – 10 min *p.i*. (**Figure 3 C)**. Under baseline conditions, the cerebellum exhibited the highest peak SUV of 3.66 at 7 min *p.i*., due to anesthesia effect on cerebellar blood flow. Peak SUVs in the brain stem, frontal cortex, globus pallidus, and occipital cortex ranged from 2.34 to 2.70. Radioactivity subsequently declined across all regions to SUVs of 0.82 – 0.99 at 120 min *p.i*. (**Figure 3 D)**. Following pretreatment with AZD9574 at both doses (0.05 mg/kg, 0.05 mg/kg), rapid uptake followed by progressive washout was also observed. Early regional SUVs were higher than at baseline, although the differences diminished later in the scan (**Figure 3 E)**.

To further quantify tracer kinetic parameters, regional TACs were fitted with 1TC and 2TC models and MA1 to obtain *V*_T_ values. In a total of 17 fits, only one 2TC *V*_T_ value demonstrated relative standard error (rSE) greater than 50% and another 2TC *V*_T_ with rSE greater than 10%, while all *V*_T_ values estimated using the 1TC model and MA1 (*t\** 30 min) were reliable. The baseline 1TC *V*_T_ values of [^18^F]Itareparib ranged from 8.5 mL⋅ cm⁻³ in the centrum semiovale to 12.5 mL⋅ cm⁻³ in the cerebellum (**Suppl. Table 1**). The *K_1_* and *k_2_* values ranged from 0.13 to 0.44 mL ⋅ cm⁻³ ⋅ min⁻¹ and 0.015 to 0.035 min⁻¹, respectively. The percentage of blocking effect was comparable using *V*_T_ values derived from all three models at a lower (1TC 21 ± 6%; 2TC 24 ± 7%; MA1 19 ± 7%) and a higher blocking dose (1TC 27 ± 3%; 2TC 31 ± 5%; MA1 26 ± 3%).

## DISCUSSION

In this study, we synthesized [^18^F]Itareparib and evaluated its imaging properties in BT142 rats and a healthy NHP under baseline and blocking conditions.

Biodistribution was first evaluated in healthy rats to characterize the pharmacokinetics of [^18^F]Itareparib. It showed high metabolic stability in the brain, with the parent tracer accounting for more than 99% of the total radioactivity. These findings indicate that the radioactivity measured in brain predominantly represented the parent tracer, with no brain penetrant radiometabolites. [^18^F]Itareparib demonstrated fast clearance from the healthy brain, while prominent liver and kidney uptake suggested hepatic and renal clearance. Tracer uptake in the spleen was reduced by 58% after AZD9574 pretreatment, consistent with the PARP1 expression in spleen and findings reported for other PARP1 PET radiotracers (*16,34–36*).

Sufficient passive brain penetration is important for the reliable PARP1 quantification across tumor regions with heterogeneous BBB integrity (*25,26,32*). To evaluate the tumor uptake and retention, dynamic [^18^F]Itareparib PET scans were performed in the rat model with patient-derived, IDH1-mutant BT142 gliomas under baseline and blocking conditions. This infiltrative glioma model was selected for its relatively preserved BBB integrity, in contrast to previously evaluated models with substantial BBB disruption (*22,26,37*), reducing the possibility of the tumor uptake being driven primarily by increased delivery through the disrupted BBB (*3,25*). Under baseline conditions, rapid initial uptake was followed by slow tumor washout and fast contralateral reference region washout, maintaining a high tumor-to-reference contrast throughout the 90-min scan. Following AZD9574 administration, tracer uptake by the tumor was reduced and activity approached contralateral reference-region levels at later time points, indicating the PARP1-specific uptake of [^18^F]Itareparib in the BT142 tumors. To establish a suitable imaging window for simplified quantification, SUVRs calculated over different time windows were compared with SRTMC DVR using contralateral brain as the reference region. SUVR_(30–60 min)_ exhibited the strongest correlation and alignment with DVR among the evaluated windows and was therefore selected as a surrogate for DVR. The significant reductions in both SUVR_(30‒60 min)_ and SRTMC DVR following AZD9574 administration indicated the PARP1-specific binding of [¹⁸F]Itareparib in BT142 tumors.

The PET findings were further supported by *in vitro* autoradiography, which demonstrated a high tumor-to-contralateral brain binding ratio and reduced tumor uptake in the presence of AZD9574. Immunofluorescence localized elevated PARP1 staining to HuNu-positive regions, with heterogeneous distributions consistent with infiltrative BT142 growth and tumor cells interspersed with normal brain tissue (*37*). Positive correlations between [^18^F]Itareparib autoradiographic signals and both PARP1 and HuNu staining further linked tracer uptake to PARP1 expression and tumor-cell distribution. Collectively, the PET imaging, autoradiography, and immunofluorescence findings support highly PARP1-specific uptake of [^18^F]Itareparib in the BT142 tumors.

NHP PET demonstrated rapid brain uptake of [^18^F]Itareparib, with cerebellum SUV greater than 3.5 at 10 min *p.i*. followed by fast washout across all regions. This reversible kinetic profile demonstrated excellent brain penetration, consistent with the contralateral brain kinetics in BT142 rats and the relatively low PARP1 expression in healthy monkey brain (*38*). Regional *V*_T_ estimates using the 1TC, 2TC, and MA1 were comparable, with consistent reductions following AZD9574 pretreatment, indicating blockable [^18^F]Itareparib binding in healthy NHP brain. Furthermore, increasing the blocking dose to 0.15 mg/kg had little influence on the magnitude of the blocking effect, which indicates nearly complete blockage at both blocking doses.

## CONCLUSION

In conclusion, [^18^F]Itareparib demonstrated fast brain penetration, high metabolic stability, and sustained tumor-to-brain contrast in BT142 rats. The reduction in tumor uptake following AZD9574 administration, supported by autoradiographic blocking and PARP1 expression in tumor tissue, indicates high PARP1-specific tracer binding in the BT142 tumors. NHP PET further demonstrated brain penetration with rapid washout from healthy NHP brain. These findings support further evaluation of [^18^F]Itareparib as a PET radioligand for PARP1 imaging in glioma and beyond.

## DISCLOSURE

This research was supported by R03CA249569, the Department of Radiology and Biomedical Imaging, and Yale School of Medicine. JC is supported by R01NS130069, R01NS123183, R41AG097311 from the National Institutes of Health. This publication was also made possible by CTSA grant UL1 RR024139 awarded jointly by the National Center for Research Resources (NCRR) and the National Center for Advancing Translational Sciences (NCATS), components of the National Institutes of Health (NIH). The contents of this publication are solely the responsibility of the authors and do not necessarily represent the official view of NIH. No other potential conflict of interest relevant to this article was reported.

## ACKNOWLEDGEMENT

We thank the staff at Yale PET Center and Yale MRRC for their technical expertise and assistance in this work.

## Key Points

### QUESTION

Can the brain-penetrant, PARP1 selective PET tracer, [^18^F]Itareparib, enable the noninvasive detection of infiltrative glioma with relatively preserved BBB integrity and the quantification of PARP1 using kinetic modeling due to its reversible kinetics?

### PERTINENT FINDINGS

In BT142 glioma-bearing rats, [^18^F]Itareparib showed approximately 4-fold greater tumor uptake than the contralateral nontumoral brain region, with tumor uptake significantly reduced by the PARP1-selective inhibitor AZD9574. Autoradiography and immunofluorescence associated tracer binding with PARP1 expression in tumor cells, while PET imaging of [^18^F]Itareparib in a healthy nonhuman primate confirmed its high brain penetration and reversible brain kinetics in healthy NHP brain, which enabled the quantitative analysis using kinetic modeling.

### IMPLICATIONS FOR PATIENT CARE

After clinical validation, [^18^F]Itareparib PET could enable more accurate delineation of brain tumors regardless their blood-brain barrier and blood-tumor barrier integrity, the noninvasive quantitative assessment of PARP1 availability for clinical occupancy studies, and the evaluation of treatment response of brain tumors.

## REFERENCES

1. Weller M, van den Bent M, Preusser M, et al. EANO guidelines on the diagnosis and treatment of diffuse gliomas of adulthood. Nat Rev Clin Oncol. 2021;18:170–186.

2. Price SJ, Jena R, Burnet NG, et al. Improved delineation of glioma margins and regions of infiltration with the use of diffusion tensor imaging: an image-guided biopsy study. AJNR Am J Neuroradiol. 2006;27:1969–1974.

3. Eidel O, Burth S, Neumann JO, et al. Tumor Infiltration in Enhancing and Non-Enhancing Parts of Glioblastoma: A Correlation with Histopathology. PLoS One. 2017;12:e0169292.

4. Law I, Albert NL, Arbizu J, et al. Joint EANM/EANO/RANO practice guidelines/SNMMI procedure standards for imaging of gliomas using PET with radiolabelled amino acids and [^18^F]FDG: version 1.0. Eur J Nucl Med Mol Imaging. 2019;46:540–557.

5. Cella E, Bosio A, Persico P, et al. PARP inhibitors in gliomas: Mechanisms of action, current trends and future perspectives. Cancer Treat Rev. 2024;131:102850.

6. Langelier M-F, Planck JL, Roy S, Pascal JM. Structural Basis for DNA Damage–Dependent Poly(ADP-ribosyl)ation by Human PARP-1. Science. 2012;336:728–732.

7. El-Khamisy SF, Masutani M, Suzuki H, Caldecott KW. A requirement for PARP-1 for the assembly or stability of XRCC1 nuclear foci at sites of oxidative DNA damage. Nucleic Acids Res. 2003;31:5526–5533.

8. Bartkova J, Hamerlik P, Stockhausen MT, et al. Replication stress and oxidative damage contribute to aberrant constitutive activation of DNA damage signalling in human gliomas. Oncogene. 2010;29:5095–5102.

9. Murnyak B, Kouhsari MC, Hershkovitch R, et al. PARP1 expression and its correlation with survival is tumour molecular subtype dependent in glioblastoma. Oncotarget. 2017;8:46348–46362.

10. Venere M, Hamerlik P, Wu Q, et al. Therapeutic targeting of constitutive PARP activation compromises stem cell phenotype and survival of glioblastoma-initiating cells. Cell Death Differ. 2014;21:258–269.

11. Galia A, Calogero AE, Condorelli R, et al. PARP-1 protein expression in glioblastoma multiforme. Eur J Histochem. 2012;56:e9.

12. Wang Y, Wild AT, Turcan S, et al. Targeting therapeutic vulnerabilities with PARP inhibition and radiation in IDH-mutant gliomas and cholangiocarcinomas. Sci Adv. 2020;6:eaaz3221.

13. Hanna C, Kurian KM, Williams K, et al. Pharmacokinetics, safety, and tolerability of olaparib and temozolomide for recurrent glioblastoma: results of the phase I OPARATIC trial. Neuro Oncol. 2020;22:1840–1850.

14. Derby S, Jackson MR, Williams K, et al. Concurrent Olaparib and Radiation Therapy in Older Patients With Newly Diagnosed Glioblastoma: The Phase 1 Dose-Escalation PARADIGM Trial. Int J Radiat Oncol Biol Phys. 2024;118:1371–1378.

15. Makvandi M, Pantel A, Schwartz L, et al. A PET imaging agent for evaluating PARP-1 expression in ovarian cancer. The Journal of Clinical Investigation. 2018;128:2116–2126.

16. Wilson TC, Xavier MA, Knight J, et al. PET Imaging of PARP Expression Using [^18^F]Olaparib. J Nucl Med. 2019;60:504–510.

17. Zhou X, Chen J, Patel JS, et al. Imaging poly(ADP-ribose) polymerase-1 (PARP1) in vivo with [^18^F]-labeled brain penetrant positron emission tomography (PET) ligand. Acta Pharmaceutica Sinica B. 2025;15:5036–5049.

18. Tong J, Chen B, Tan PW, Kurpiewski S, Cai Z. Poly (ADP-ribose) polymerases as PET imaging targets for central nervous system diseases. Front Med (Lausanne). 2022;9:1062432.

19. Tong J, Chen B, Volpi T, Li Y, Ellison Paul A, Cai Z. Current Advances in PARP1-Targeted Theranostics. Journal of Labelled Compounds and Radiopharmaceuticals. 2025;68:e4135.

20. Carlucci G, Carney B, Brand C, et al. Dual-Modality Optical/PET Imaging of PARP1 in Glioblastoma. Mol Imaging Biol. 2015;17:848–855.

21. Young RJ, Demetrio De Souza Franca P, Pirovano G, et al. Preclinical and first-in-human-brain-cancer applications of [^18^F]poly (ADP-ribose) polymerase inhibitor PET/MR. Neurooncol Adv. 2020;2:vdaa119.

22. Carney B, Carlucci G, Salinas B, et al. Non-invasive PET Imaging of PARP1 Expression in Glioblastoma Models. Mol Imaging Biol. 2016;18:386–392.

23. Chan CY, Hopkins SL, Guibbal F, et al. Correlation between molar activity, injection mass and uptake of the PARP targeting radiotracer [^18^F]olaparib in mouse models of glioma. EJNMMI Res. 2022;12:67.

24. Cselényi Z, Jucaite A, Stragliotto G, et al. Subtype-selective positron emission tomography imaging of poly ADP-ribose polymerase-1 binding in non-human primates, healthy volunteers, and patients with glioblastoma using [^11^C]AZ14193391. Neuro Oncol. 2026;28:802–814.

25. Kizilbash SH, Gupta SK, Chang K, et al. Restricted Delivery of Talazoparib Across the Blood-Brain Barrier Limits the Sensitizing Effects of PARP Inhibition on Temozolomide Therapy in Glioblastoma. Mol Cancer Ther. 2017;16:2735–2746.

26. Chen B, Ojha DP, Toyonaga T, et al. Preclinical evaluation of a brain penetrant PARP PET imaging probe in rat glioblastoma and nonhuman primates. Eur J Nucl Med Mol Imaging. 2023;50:2081–2099.

27. Reilly SW, Puentes LN, Schmitz A, et al. Synthesis and evaluation of an AZD2461 [^18^F]PET probe in non-human primates reveals the PARP-1 inhibitor to be non-blood-brain barrier penetrant. Bioorg Chem. 2019;83:242–249.

28. Montagnoli A, Papeo G, Rainoldi S, et al. Abstract 4843: NMS-P293, a PARP-1 selective inhibitor with no trapping activity and high CNS penetration, possesses potent in vivo efficacy and represents a novel therapeutic option for brain localized metastases and glioblastoma. Cancer Research. 2018;78:4843–4843.

29. Wilson AA, Jin L, Garcia A, DaSilva JN, Houle S. An admonition when measuring the lipophilicity of radiotracers using counting techniques. Appl Radiat Isot. 2001;54:203–208.

30. De Feyter HM, Behar KL, Rao JU, et al. A ketogenic diet increases transport and oxidation of ketone bodies in RG2 and 9L gliomas without affecting tumor growth. Neuro Oncol. 2016;18:1079–1087.

31. Staniszewska AD, Pilger D, Gill SJ, et al. Preclinical Characterization of AZD9574, a Blood-Brain Barrier Penetrant Inhibitor of PARP1. Clin Cancer Res. 2024;30:1338–1351.

32. Zhou X, Chen J, Patel JS, et al. Imaging poly(ADP-ribose) polymerase-1 (PARP1) in vivo with [^18^F]-labeled brain penetrant positron emission tomography (PET) ligand. Acta Pharm Sin B. 2025;15:5036–5049.

33. Waterhouse RN. Determination of lipophilicity and its use as a predictor of blood-brain barrier penetration of molecular imaging agents. Mol Imaging Biol. 2003;5:376–389.

34. Michel LS, Dyroff S, Brooks FJ, et al. PET of Poly (ADP-Ribose) Polymerase Activity in Cancer: Preclinical Assessment and First In-Human Studies. Radiology. 2017;282:453–463.

35. Guibbal F, Hopkins SL, Pacelli A, et al. [^18^F]AZD2461, an Insight on Difference in PARP Binding Profiles for DNA Damage Response PET Imaging. Mol Imaging Biol. 2020;22:1226–1234.

36. Tornblom A, He C, Bolin M, et al. Biodistribution and dosimetry of the carbon-11 labelled PARP1-selective PET radiopharmaceutical [^11^C]AZ14193391. EJNMMI Res. 2026;16.

37. Luchman HA, Stechishin OD, Dang NH, et al. An in vivo patient-derived model of endogenous IDH1-mutant glioma. Neuro Oncol. 2012;14:184–191.

38. Ferreira MT, Berger L, Rouleau M, Poirier GG. Assessment of PARP-1 Distribution in Tissues of Cynomolgus Monkeys. J Histochem Cytochem. 2020;68:413–435.

